# LoopViz: A uLoop Assembly Clone Verification Tool for Nanopore Sequencing Reads

**DOI:** 10.1101/2021.02.01.427927

**Authors:** Mariele Lensink, Bradley W. Abramson, Nolan Hartwick, Alfonzo Poire, Vincent A. Bielinski, Todd P. Michael

**Affiliations:** Plant Molecular and Cellular Biology, Salk Institute of Biological Sciences, La Jolla, CA, USA; J. Craig Venter Institute, La Jolla, CA, USA

## Abstract

Cloning has been an integral part of most laboratory research questions and continues to be an essential tool in defining the genetic elements determining life. Cloning can be difficult and time consuming as each plasmid is unique to a particular project and each sequence must be carefully selected, cloned and sequenced to determine correctness. Loop assembly (uLOOP) is a recursive, Golden Gate-like assembly method that allows rapid cloning of domesticated DNA fragments to robustly refactor novel pathways. With uLOOP methodologies, one can clone several sequences directionally to generate a library of transcriptional units (TUs) in plasmids within a single reaction but analysis of the plasmid population has been impeded by current sequencing and analysis methods. Here we develop LoopViz, a quality control tool that quantifies and visualizes results from assembly reactions using long-read Oxford Nanopore Technologies (ONT) sequencing. LoopViz identifies full length reads originating from a single plasmid in the population, and visualizes them in terms of a user input DNA fragments file, and provides QC statistics. This methodology enables validation and analysis of cloning and sequencing reactions in less than a day, determination of the entire plasmid’s sequence, and sequencing through repetitive meta-regions that cannot be meaningfully assembled. Finally, LoopViz represents a new paradigm in determining plasmid sequences that is rapid, cost-effective and performed in-lab. LoopViz is made publicly available at https://gitlab.com/marielelensink325/loopseq

## 1 Introduction

Cloning has been an integral part of any laboratory research question and continues to be an essential tool in defining the genetic elements determining life. Complementation assays in bacterial gene knockout lines, integration of plant T-DNA cassettes and CRISPR/Cas genome editing would not be possible without the ability to clone and determine the sequences of plasmids. Almost all cloned sequences are validated with some type of sequencing and usually only the region of interest is sequenced due to the cost and labor involved in designing, pipetting and sequencing short stretches (<1kb) of sequence using Sanger sequencing. Next generation sequencing (NGS) can be used if whole plasmid sequences are needed but can be costly or excessive if only one plasmid requires sequencing (i.e. Illumina). Additionally, short read sequencing often lacks the contiguity needed for larger plasmids or populations of plasmids that may contain stretches of repetitive DNA (i.e. CRISPR arrays) (Kunin, Sorek, and Hugenholtz 2007).

Cloning has become increasingly more advanced as researchers develop new methods from restriction enzymes to homologous recombination based cloning (e.g. Gibson assembly; Gateway) (Gibson et al., n.d.) to TypeIIs assembly methods (e.g. Golden Gate; uLOOP) (Chao, Yuan, and Zhao 2015). uLoop assembly is a type IIs assembly method that utilizes SapI and BsaI restriction enzymes for combinatorial and iterative assembly of domesticated synthetic DNA fragments, hereafter synDNA, into domesticated receiving vectors already compatible with transformation into bacteria, diatoms or plants (Pollak et al. 2020). Following uLOOPs standard syntax the SapI and BsaI sites are removed and create compatible “sticky-ends” for ligating fragments into a domesticated level-0 vector. The recursive nature allows four level-0 parts to be cloned into a level-1 vector, four level-1 parts into level-2 vectors and then four level-2 parts back to level-1 vector and so on. One can assemble a library or population of plasmids if multiple synDNA fragments with the same overlapping “sticky-ends” are added to one digestion/ligation reaction. As gene synthesis costs decrease researchers can effectively synthesize DNA fragments (synDNA) for cloning with the added benefit of codon optimization or removal of difficult to clone hairpin structures. This makes cloning relatively rapid in the lab where a single researcher is limited by cost and design time generally.

Currently, the standard method for validating a successfully cloned sequence utilizes sanger sequencing, which can be slow and unlikely to produce comprehensive results for combinatorial cloned uLoop reactions. uLoop reactions can generate a population of plasmids featuring random combinations of desired synDNA layouts, and successful Sanger sequencing would first require isolating individual plasmids from that population since identical sequences in identical fragments could occur in multiple assembled plasmids. If the synBlocks are sufficiently large it may require multiple Sanger sequencing runs to determine, for that plasmid, which combination of synBlocks were inserted. Single molecule long read sequencing technologies offer an alternative that can sequence directly on the reaction products of a combinatorial cloning reaction.

In this study, we show that detection of full-length plasmid sequences can be determined on a population of plasmids cloned using uLoop assembly. We generated several cloning reactions and sequenced the ligation reaction directly, a collection of 5 purified plasmids and finally an entire population of cloned plasmids. We designed LoopViz to determine full length plasmids in an ONT sequencing dataset and define the DNA elements within the full length plasmid reads. LoopViz then generates graphs to visualize the data. The approach outlined here represents an entirely new method for validating cloned sequences and using LoopViz for visualizing sequencing data of cloning reactions.

## 2 Materials and Methods

LoopViz implements a snakemake workflow system for easy modification and versatile analysis, and it is run from the command line. It is written mostly in python and utilizes packages such as pandas and bokeh for data formatting and visualization.

The same biological assembly reaction was set up using loop assembly methods and “synBlocks,” or synDNA fragments used assembled together in the cloning reaction, as insertion components. Reaction products were harvested at three different times that were hypothesized to change the composition of the number of full-length reads and number of unique plasmids in the population. This data was used to analyze the performance and accuracy of our tool, LoopViz.

### 2.1 Program Development

LoopViz determines full length plasmids by analyzing the composition of inserted components, or “synBlocks” via user defined settings such as the number of insertion positions as well as what components should be present. LoopViz does this by using Blastn to annotate fastq raw read output using standard backbone and selectable marker sequences provided internally as well as a user provided database of all possible insertion components. The program then runs an overlap analysis and quality control of each annotated read and filters for full length plasmid sequences. After annotating and determining full length plasmids, LoopViz then provides a diverse set of output files for a wide array of analysis.

#### 2.1.1 Determining Full Length Reads

Because there is great variation in the size of inserted components, it is impossible to detect a full plasmid based on length when multiple plasmids with different lengths are sequenced together. Furthermore, the composition of reads can be altered by sequencing a ligation reaction high in unassembled fragments versus a plasmid isolated from a transformed bacterial colony which should have few fragments and only one plasmid. Since each synDNA fragment is of different length and the composition of each plasmid is mostly random, the exact length of the sequencing read for any full length plasmid is varied. For this reason, we chose to determine a full length plasmid reads by analyzing the presence and composition of its components. LoopViz contains two databases of standard loop assembly parts (backbone sequences and selectable marker genes) that it uses to annotate each read. For convenience, the standardized internal databases are separated into two by sequence type for easy substitutions should the user desire to alter their experiment more. It also requires a database of insertion components from the user as well as the number of insertion positions that should be present on a full length plasmid. In this way, we have all possible parts of a full length plasmid used in the reaction and are subsequently able to determine if it is full length by confirming all necessary parts are present.

#### 2.1.2 Annotation of Plasmid Components

We used NCBI’s blastn to annotate synBlocks, backbone genes, and selectable markers onto our fastq sequences and generate a customized table output containing information on each alignment (figure 1a). Blast hits must have a minimum of 70% coverage of the specific synBlock to be included in the blast output. Three separate blast tables are generated from this step, one for synBlock hits, and two for the internal inputs (backbone genes and selectable marker sequences) and added to an output directory generated by the program.

**[Figure 1a].**
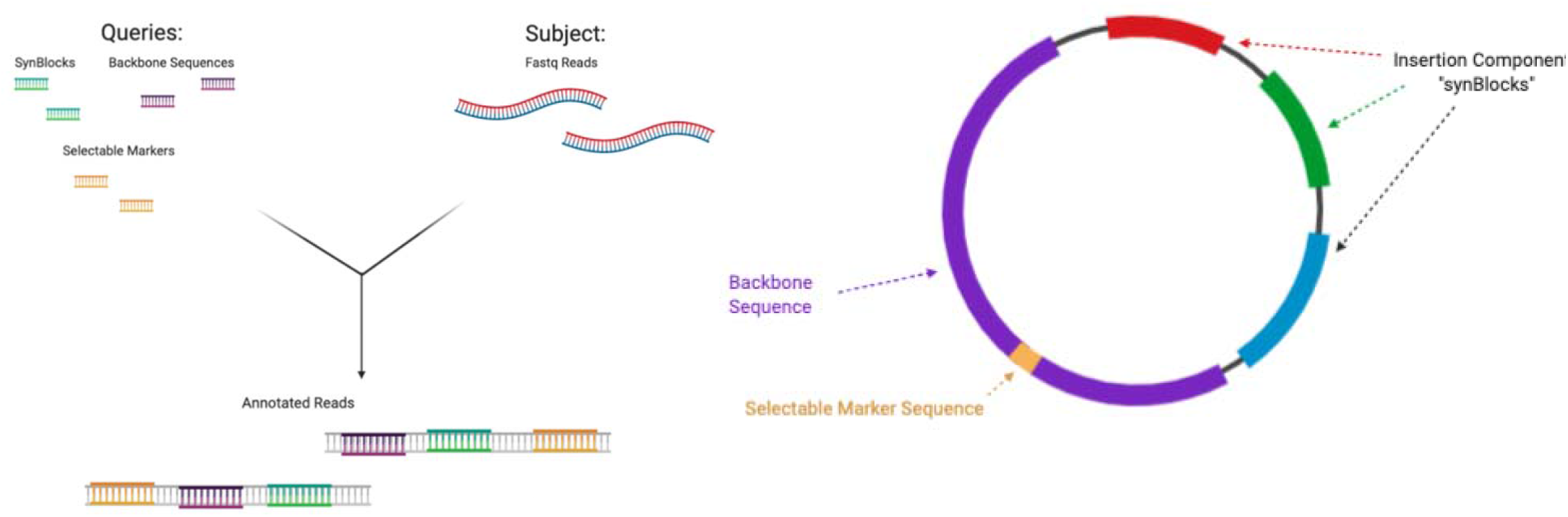
Layout of annotation step. Each of the 3 databases are used as queries to annotate the user inputed fastq sequence file. The output of this step is a series of blast tables of every annotation found. The tables are then filtered and reformatted to represent single reads and each of the quality controlled annotations on them. [Figure 1b] Layout of a hypothetical full length plasmid containing all necessary components annotated, cleaned, identified, and later analyzed by LoopViz. Other reads that do not contain all necessary parts displayed in Figure 1b are removed from analysis. (Created using Biorender.com)

It is important to note that LoopViz does not support synBlocks under 30 nucleotides in length. This is the result of both a software limitation and a sequencing limitation of ONT. Blastn will not align sequences under 30 nucleotides in length using the standard algorithm. While Blastn features settings for short sequences, doing so dramatically reduces the sensitivity of Blastn in combination with the high error rates in Nanopore sequencing reads. A work around for this limitation is to combine the synBlock position featuring a short synBlock with a neighboring synBlock position.

#### 2.1.3 Overlap analysis

After the reads have been annotated and blast tables have been generated, they must be cleaned using an overlap analysis. Before we can determine that a read has the correct number of insertion components, we must first remove any incorrect annotations. If there are two synBlocks present with an overlap greater than 30 nucleotides, the synBlock with a lower percent identity is removed from the annotations(Figure 2). The overlap cleaned synBlock alignments are then passed on for subsequent read level filtering.

**[Figure 2].**
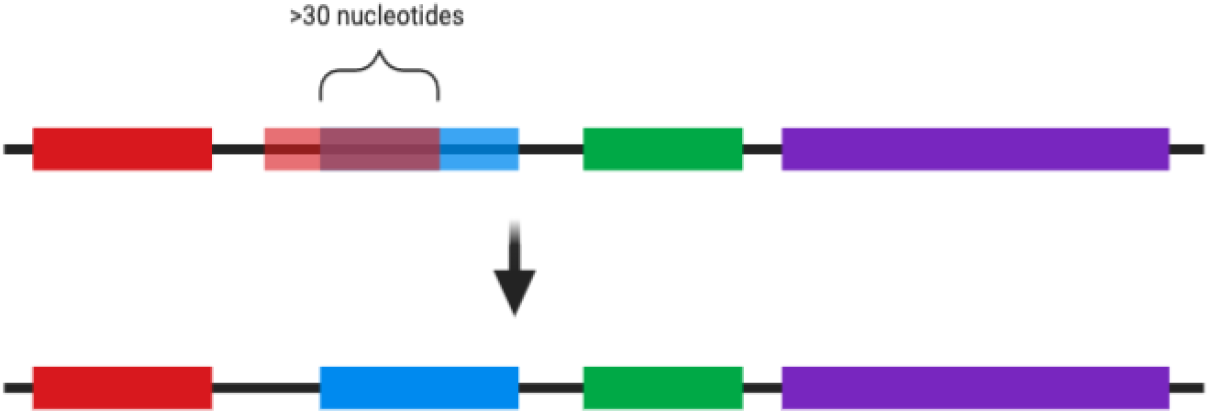
Diagram of overlap analysis displaying h overlaps are detected and removed. When there is an overlap between two synBLocks larger than 30bps, the synBlock with a lower percent identity value is removed. (Created using Biorender.com)

#### 2.1.4 Filtering

Each type of blast annotation table has its own set of filtering criteria unique to the type of annotation. A full-length reads must contain a backbone sequence, at least one selectable marker sequence, and the correct number of inserted components as defined by the user, in this case three synBlocks. Reads that do not meet the criteria for all three filters are deemed incomplete and removed from the analysis. Reads passing all three criteria are considered full length.

#### 2.1.5 Read formatting for analysis

After filtering, the reads are formatted into a simplified representation purely in terms of synBlocks and a single consistent backbone element (pMB1_ori). Because the sequenced read could start at any location on the circular plasmid, the read with all of its annotated synBlocks, is rotated to a common starting position, specifically the pMB1_ori. Reads can also be sequenced on either the template or the coding strand of the DNA. In response to this, each read with a “minus” orientation for its determining backbone element (pMB1_ori) is reverse complemented. This leaves us with a very simple representation of the full length reads, containing only synBlocks, their location, and orientation -all relative to the plasmid backbone sequence.

#### 2.1.6 output files

LoopViz creates an output directory with many different files providing a wide variety of information for analysis by the user (Table 1).

**[Table 1].** Table displaying file names and brief descriptions of relevant output files generated by LoopViz and located in the LoopViz output directory.

| File Name | Description |
| --- | --- |
| Loopseq_viz.html | An html page containing interactive graphics displaying the composition and frequency of full length plasmid |
| Plasmid_types.counts.tsv | A list of unique plasmid types and their frequency. Formatted as a table with each insertion component listed per column and the number of times that specific plasmid was found in the final column |
| full_length_reads.fasta | FastA file containing all of the filtered, full length plasmid sequences |
| Read_types.tsv | A table containing the sequence ID of each full length read and its corresponding insertion component annotations |
| SummaryStats.tsv | A table containing statistics/quantitative information on read output available for further analysis |
| *blast.tbl | Three tables from blast - one for backbones, selectable markers, and insertion components - containing annotations for each read as well as supplemental information on each blast hit (description below) |

### 2.2 Experimental set up

Here we use a type-II restriction enzyme cloning system(uLoop) to clone multiple fragments in one assembly reaction. This type of cloning produces a large number of unique plasmids that can be difficult to analyze effectively with current sequencing technologies. Briefly, a ligation reaction, plasmids isolated from picked transformed colonies and a population of plasmids isolated from transformed E. coli carrying the cloned library were sequenced using Nanopore long read sequencing for analysis using LoopViz.

#### 2.2.1 uLoop assembly

uLoop assemblies were generated following uLoop protocols as previously described (Pollak et al. 2020); dx.doi.org/10.17504/protocols.io.yxnfxme. Briefly, synBlocks were ordered from IDT with appropriate BsaI and SapI overhangs for assembly. Each cloning reaction contained a total of 19 fragments where 5 AC promoters, 11 CD CDS, 1 DE 3xStop and 2 EF terminators and a Level-1 vector (S Table 1). These were assembled for 25 cycles and samples were taken after ligation and used for transformation into competent cells (NEB:C2987I). The transformation reaction was split in two and plated on selective LB (Spec 50mg/L) or selected in LB liquid media with spec and selection was performed overnight at 37C (Figure 3). This experiment was performed 3 separate times resulting in 9 samples for sequencing. Five plasmids from single colonies or from a population of *E. coli* carrying unique plasmids grown in liquid cultures were extracted using Qiagen’s miniprep spin columns (#27104).

**[Figure 3].**
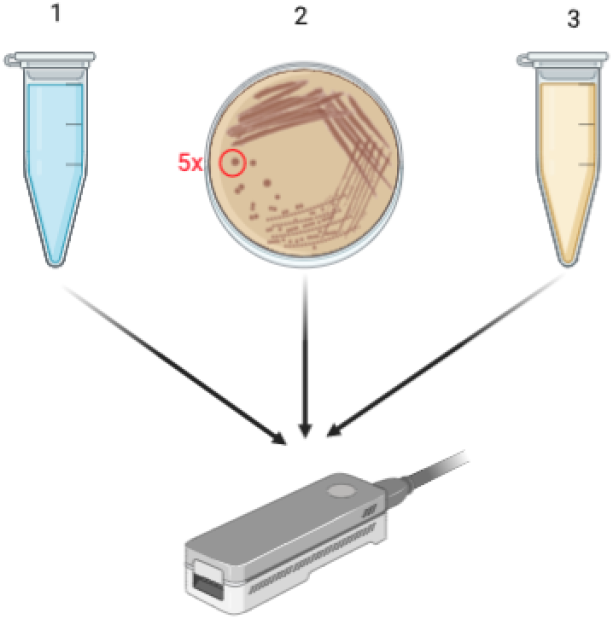
Layout of experimental setup. uLoop reaction was performed in triplicate and products were harvested using 3 different methods before sequencing. Extraction method 1: DNA products were taken directly from the uLoop reaction to sequencing. Method 2: Products were first transformed into bacteria via heat shock, plated, and extracted from 5 harvested colonies. Method 3: Products were transformed into liquid culture and extracted from the total population of cells. All extracted plasmid samples were then library prepped and sequenced using an Oxford Nanopore GridION. Raw fastq files generated by sequencing were then used as input for LoopViz. (Created using Biorender.com)

#### 2.2.2 Sequencing

uLoop ligations or isolated plasmids were used as DNA inputs into Nanopores’s rapid barcoding kit (SQK-RBK004). Each sample (7.5ul), ligation reaction, 5 plasmid mixture or population of plasmids was barcoded with 1ul RAP and sequenced on one flongle flowcell run on a GridION machine. Reads were basecalled with the MinKnow software running guppy v3.2 in high-accuracy settings. The uLOOP cloning, barcoding and sequencing was performed in triplicate.

## 3 Results

Cloning with uLOOP assembly is relatively rapid and numerous constructs can be cloned in a single day with a high degree of accuracy yet clone verification can take multiple days. Initially we designed and synthesized DNA (SynDNA) fragments with uLOOP compatible type IIs restriction overhangs to clone complete transcriptional units into receiver vectors (S Table 4). The SynDNA was designed such that a plant promoter and 5’UTR had AC overhangs, coding sequences had CD overhangs, a short 3xSTOP fragment contained DE overhangs and 3’UTR’s contained EF overhangs to clone into the AF receiver plasmid following standard uLOOP assembly methods and nomenclature (S Table 4). A total of five AC promoters, nine CD coding sequences, one DE 3xSTOP and two EF terminators were diluted to equal molar concentrations and placed into a single uLOOP digestion/ligation reaction. Following thermocycling digestion/ligation an aliquot was taken for sequencing and the rest of the reaction was transformed into Dh5a competent *E. Coli*. Part of the recovered transformed cells were plated on selective LB agar plates and the rest of the reaction was grown in selective LB liquid media overnight. The following day, 5 independent colonies were picked from the LB plate, selected overnight and plasmids were extracted independently. A plasmid miniprep was also performed on the antibiotic selected, liquid population of transformants (Figure 3).

Of the output provided by LoopViz, most notable is the loopseq_vis.html file, which provides graphic visualizations of the uLoop reaction results(figure 4). The html file first displays a pie chart representing the composition of reads categorized by unique plasmid types (figure 4a). Plasmids with identical compositions of synBlocks are clustered into a “unique plasmid type” group. The number of plasmids in each unique plasmid type group are then represented in the pie chart as percentages of the whole read population. The other feature is a multiplex of legends that correspond to a heat map representing the entirety of whole length reads and their corresponding synBlocks (Figure 4b). The number of legends displayed corresponds to the number of insertion positions in the uLoop reaction, specified by the user in the command that executes the program. The number of different colors present for each position’s legend corresponds to the number of different synBlocks detected by LoopViz from the reaction’s full length reads. For example, in the legend representing synBlock position 3 (figure 4b), there are only two options for synBlocks -tNOS terminator and tHSP terminator -because only these two synBlocks were incorporated into the uLoop reaction for position 3. The legends describe the combination of synBlocks for each read in the corresponding heatmap (figure 4c). In the heatmap, each row represents a single full length plasmid, and each column represents a synBlock insertion position. Each detected synBlock in the reaction is assigned a color and represented by that color when it is present on a read in the heatmap. The heatmap represents both a general composition of unique types, similar to that of the pie chart, as well as a more in depth display of the composition and frequency of synBlocks present in the reaction results.

**[Figure 4].**
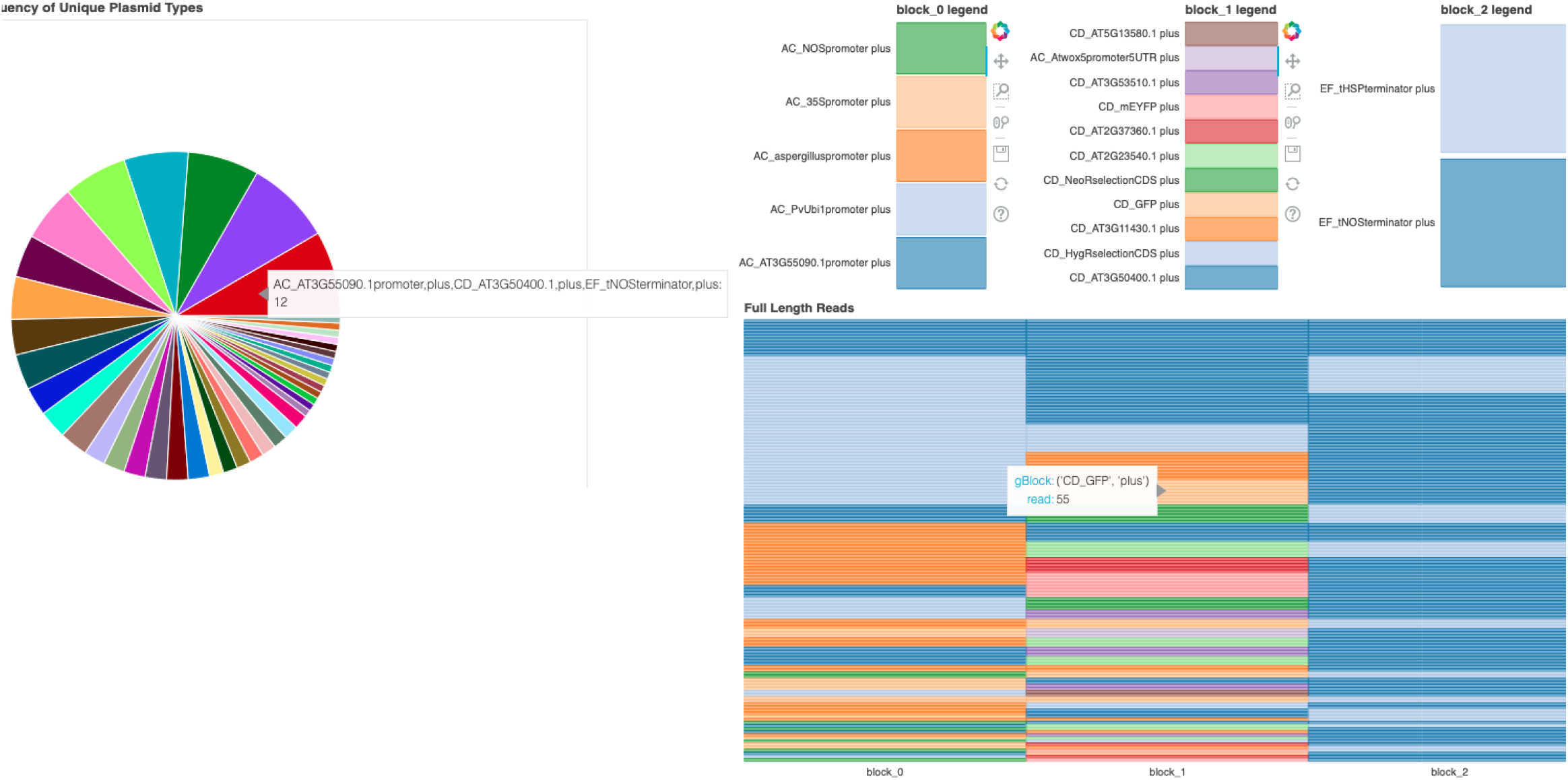
Example output from loopseq_vis.html a) Pie chart displaying the proportion of unique plasmid types (determined by combination of inserted sequences). A hover tool is included in place of a legend for a more condensed display of information. b) Heatmap of full length read combinations with accompanying legend. There is a separate legend for each insertion position. Heat map displays each full length read and it’s corresponding synBlocks. Each column represents one of the inserted sequences on that read (in order). The number of columns will correspond with the number of insertion positions determined by the user.

### 4.2 performance on uLoop combinatorial experiment

Experiments where plasmids were transformed (extraction methods 2 and 3) had greater average sequence length in their original fastq output than experiments sequenced directly from the Uloop reaction (figure 3, figure 5a). This supports our hypothesis that transforming the reaction sequence into E. coli would select for full length pieces and subsequently increase the average length of reads sequenced. The average length of reads was highest in method 3, total population in liquid broth, with an average length ranging from 3,311 to 3,479 bp. Average length was lowest in method 1, reaction fragments direct from the experiment, and ranged between 651 and 799 bp. Plasmids from plated colonies (method 2) were also in larger quantities than those directly extracted from the reaction (method 1), but smaller in length compared to method 3 (liquid population), generating questions on the effect of selecting colonies as opposed to extracting from a suspended liquid population. It is likely that the total number of full length reads is highest in the sequenced plasmid datasets and not the ligation dataset since the uLOOP ligation reaction still contains the small fragments used for cloning (figure 5b). Both reaction replicates 2 and 3 indicate that method 3 (liquid population) contains the highest number of full length reads with values of 101 and 143 reads, respectively, followed by plasmids transformed into plated colonies with 52 and 13 full length reads. This relationship is inverted, however, in the first reaction replicate. In all 3 reaction replicates, method 1 (assembly reaction products) contained the lowest number of full length reads, ranging from 1 to 5 reads detected. The highest number of full length reads detected by LoopViz was from the liquid population (method 3) in reaction replicate 3 with 143 reads.

**[Figure 5].**
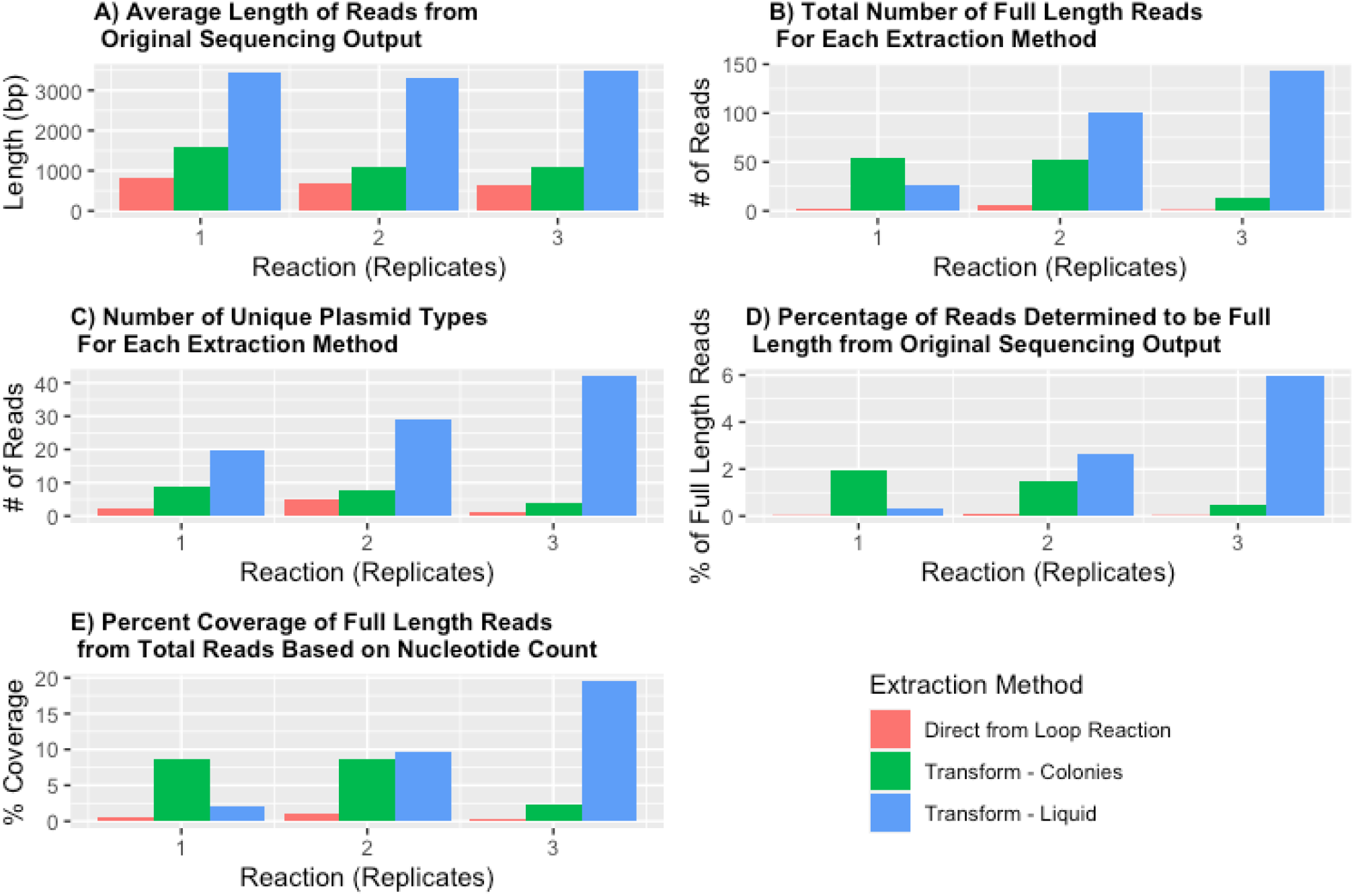
Experiment was run in triplicate, using three separate uLoop assembly reactions. Plasmids were extracted from each replicate using 3 different methods. In method 1, reaction fragments were directly sequenced. In method 2, reaction fragments were transformed into E. coli to select for whole plasmids and then plated. Plasmids were then extracted from 5 selected colonies. For method 3, reaction fragments were transformed into E. coli to select for whole plasmids. The E. coli were then grown in liquid broth and plasmids were extracted from the total population of cells. a)Displays the average length of reads from the original fastq sequence file. b) Bar graph displays the total number of full length reads filtered for each extraction method grouped by reaction replicate. c) Bar graph showing the number of unique plasmid types identified by LoopViz for each extraction method. d) Displays percentage of reads from original sequencing output that were determined to be full length. e) Displays coverage from full length reads over the original sequencing output file calculated by base pair count.

For each reaction replicate, method 3 contained the greatest diversity of plasmids determined by the number of unique full length plasmid types detected by LoopViz, and ranged from 20 to 42 unique plasmids (6c). This is expected, as the method extracts from a population of transformed *E. coli* containing many plasmids. In contrast, method 1 contained the lowest number of unique plasmid types, ranging from 1 to 5. This result is likely because most of the sequencing reads were determined to be individual synBlock fragments. There was significant variation in the number of unique plasmid types between the 3 methods, with an Anova p value of .0047. A tukey HSD test was then run and determined that significant variation was between method 3 and method 1 (p value 0.0054) as well as method 3 and method 2 (p value (0.012). These results follow a similar trend to that of figure 5a displaying average read length amongst the 3 different extraction methods (fig 4) in each reaction. These results also support our hypothesis that extracting from a population of plasmids suspended in liquid broth would contain a greater number of unique plasmid types than selecting from 5 picked colonies on an agar plate. It should also be noted that more than 5 unique plasmids were detected for reactions 1 and 2 even though only 5 colonies were selected, possibly indicating that this particular lab strain of E. coli transformed more than 1 plasmid per cell or that there were more colonies accidentally picked (6c).

The total percentage of full length reads from the total sequencing output detected by LoopViz never exceeded 6%. Method 1 had the lowest percentage of full length reads from the total in all 3 replicates (never exceeding 0.095%), which is consistent with our hypotheses. For reaction replicates 2 and 3, method 3 had the highest percentage of full length reads followed by method 2, however this relationship is reversed in reaction replicate 1 -a similar trend to that found in figure 5b.

The Full Length Read Nucleotide Fraction, meaning the total length of full length reads divided by the total length of reads for a sample, displays similar trends to those of figures 6a and 6d (figure 5e), with method 3 in reaction replicate 1 being an outlier. The highest percent coverage was 19.5% in reaction replicate 3 for method 3.

## 5 Discussion

Here we used Nanopore’s rapid transposon based barcoding of plasmids for sequencing to determine orientation and composition of combinatorial cloning reactions and developed a tool, LoopViz, to detect full length plasmids in the sequenced dataset. We generated uLOOP assembly cloning (Pollak et al. 2020) reactions and sequenced them on the third-generation long read Nanopore flongle flow cell to determine how the cloning reactions were put together. We were able to detect and filter full length reads from fragmented reads with a newly designed program, LoopViz, which detects full length reads and determines the order, direction and identity of the cloned insertion sequences. This represents the first time to our knowledge that populations of combinatorial cloned plasmids have been sequenced on Nanopore’s flongle flow cell. Prior to the development of LoopViz, analysis of these sequencing products would have been impossible due to the highly similar and numerous generated plasmids in a single reaction. By exploiting the long read potential of Nanopore sequencing, LoopViz is able to accurately determine the composition of a large combinatorial cloned library.

One limitation to combinatorial cloning experiments is the inability to determine the composition of the sequences that were cloned in a population due to short read lengths with common sequencing technologies (i.e. sanger sequencing). Sanger sequencing requires that plasmids in a population have unique sequences so unique sequencing primers can be designed to determine the sequence of a roughly 1kb region within the plasmid. With third generation sequencing technologies the limitation to sequencing length is often the ability to extract high molecular weight (HMW) DNA and since plasmids are generally small <20kb we used Nanopore to sequence full length plasmids that can be observed and analyzed as a single read. This type of sequencing can be performed in any lab using Nanopore’s flongle flowcell which can be run on a laptop in an afternoon.

Sequencing plasmids with Nanopore’s flongle flow cells can be performed at a greatly reduced cost compared to Sanger sequencing and represents a potentially new paradigm in sequencing plasmids. The cost benefits of nanopore-based sequencing of cloning reactions allow for scientists to perform in depth quality control on their experiments, even as cloning products increase in size and complexity. With the potential release of Nanopore’s 96 rapid barcoding kit, 96 plasmids could be sequenced on a single flongle flowcell for a cost of roughly 0.39$ /kb compared with 7$ /kb with Sanger sequencing (S table 3). Despite the advantages of sequencing plasmids with Nanopore, there are still complications in library prep and sequencing affecting its performance. The major limitation currently is the per read error in sequencing and is a major limiting factor. In the future, the release of new pore types may increase read accuracy thereby allowing for greater adoption of these protocols (Tytgat et al. 2020).

Currently, the percentage of reads determined to be full length is low. This could be attributed to poor optimization in the biological reaction and/or the filtering method in the LoopViz algorithm. The molar ratio of the amount of transposase for barcoding to the number of plasmids in the reaction directly impacts the number of full length reads by changing the expected number of times each plasmid is cut. If too much transposase is used, then many of the plasmids will be cut multiple times, and very few of the sequenced reads will represent complete plasmids. If less transposase is used then the likelihood that only one plasmid receives a single barcode increases thereby increasing the number of full length reads representing full-length plasmids. Careful optimization of the molar ratio will be a requirement for future reactions as opposed to the methodology used here where sample volumes were kept constant and the molar ratio was not determined.

Additionally, for this particular experiment, LoopViz required exactly three synBlocks to identify a read as full length. If synBlocks are not detected in the read accurately then the criteria of full-length reads cannot be satisfied and actual full length reads would not be detected simply because they did not contain three annotated synBlocks. Here we use blastn to find alignments that have 70% coverage over the synBlock fragment region of the read, which is appropriate for the expected Nanopore error rate. However, an alternative aligner with better optimized alignment parameters like Minimap2 may offer further improvement in detection of synBlocks within noisy plasmid reads, thereby increasing the accuracy of block annotation and the percent of full-length reads detected.

