## Supplemental Figures for "LoopViz: A uLoop Assembly Clone Verification Tool for Nanopore Sequencing Reads"

### Supplemental Information

| Column | Content |
|---|---|
| 1 | Read ID (Subject) |
| 2 | Inserted Sequence (Query) |
| 3 | Subject Strand |
| 4 | Subject Length |
| 5 | Subject Start |
| 6 | Subject End |
| 7 | Query Length |
| 8 | Query Start |
| 9 | Query End |
| 10 | ALignment Length |
| 11 | E Value |
| 12 | Percent Identity |

[Supplemental Table 1] Description of blast table columns

| Reaction | Barcode | FullLengthReads | TotalReads | UniqeCount | FullReadsNucLength | TotalReadsNuclength | NucLengthPercent | FullAvgLength | TotalAvgLength | FullLengthPercent |
|---|---|---|---|---|---|---|---|---|---|---|
| 1 | 1 | 2 | 8103 | 2 | 29977 | 6474649 | 0.4629903 | 14988.500 | 799.0434 | 0.02468222 |
| 1 | 2 | 55 | 2868 | 9 | 390754 | 4549006 | 8.5898766 | 7104.618 | 1586.1248 | 1.91771269 |
| 1 | 3 | 26 | 7348 | 20 | 510121 | 25405808 | 2.0078913 | 19620.038 | 3457.5133 | 0.35383778 |
| 2 | 1 | 5 | 5264 | 5 | 43012 | 3698847 | 1.1628489 | 8602.400 | 702.6685 | 0.09498480 |
| 2 | 2 | 52 | 3560 | 8 | 338998 | 3966105 | 8.5473783 | 6519.192 | 1114.0744 | 1.46067416 |
| 2 | 3 | 101 | 3853 | 29 | 1235721 | 12758236 | 9.6856728 | 12234.861 | 3311.2473 | 2.62133403 |
| 3 | 1 | 1 | 3934 | 1 | 7219 | 2561260 | 0.2818535 | 7219.000 | 651.0574 | 0.02541942 |
| 3 | 2 | 13 | 2906 | 4 | 71979 | 3121983 | 2.3055539 | 5536.846 | 1074.3231 | 0.44735031 |
| 3 | 3 | 143 | 2403 | 42 | 1630018 | 8360385 | 19.4969251 | 11398.727 | 3479.1448 | 5.95089471 |

[Supplemental Table 2] Table of values used in figure 5a-e

| cost in $USD | Nanopore | Sanger |
|---|---|---|
| RBK-004 kit | 650 | 0 |
| Primer | 0 | 3.5 |
| RXN (Retrogen) | 0 | 3.5 |
| Flowcell cost | 90 | 0 |
| Length (kb) | 20 | 1 |
| Samples multiplexed | 96 | 1 |
| Total cost | 0.39 | 7.00 |

[Supplemental Table 3] costs associated with sequencing plasmids. Sanger(Retrogen/Eaton) and Nanopore costs were determined from the website listed price.

| | | |
|---|---|---|
| SynBlock | CD_AT2G23540.1 | GGTCTC |
| SynBlock | CD_AT2G37360.1 | GGTCTC |
| SynBlock | CD_AT3G11430.1 | GGTCTC |
| SynBlock | CD_AT3G50400.1 | GGTCTC |
| SynBlock | CD_AT3G53510.1 | GGTCTC |
| SynBlock | CD_AT5G13580.1 | GGTCTC |
| SynBlock | CD_HygRselectionCDS | GGTCTC |
| SynBlock | CD_NeoRselectionCDS | GGTCTC |
| SynBlock | EF_tNOSterminator | GGTCTC |
| SynBlock | AC_aspergilluspromote | GGTCTC |
| SynBlock | AC_35Spromoter | GGTCTC |
| SynBlock | EF_tHSPterminator | GGTCTC |
| SynBlock | AC_PvUbi1promoter | GGTCTC |
| SynBlock | AC_NOSpromoter | GGTCTC |
| SynBlock | AC_AT3G55090.1prom | GGTCTC |
| SynBlock | AC_Atwox5promoter5U | gtctcAGG |
| SynBlock | CD_GFP | gtctcAGG |
| SynBlock | CD_mEYFP | gtctcAGG |
| BB_components | UNSX | ccaggata |
| BB_components | UNS1 | cattactcg |
| BB_components | UNSp2 | GCTGGGA |
| BB_components | UNSp3 | GAGCCAA |
| BB_components | UNSpX | CTACAAC |
| BB_components | RB_overdrive | ccttgacagg |
| BB_components | LB | GATCTTGG |
| BB_components | lacZ_alpha | ATGACCA |
| BB_components | UNStoUNS_731 | CATTACTC |
| BB_components | pMB1_ori | tttccatagg |
| BB_components | p0dd_backbone | CATTACTC |
| BB_components | pCAe_bb | cacagaatc |
| BB_components | pCAo_bb | CATTACTC |
| BB_components | pEven_bb | CATTACTC |
| BB_components | pGreenII_bb | TTTTTATC |
| BB_components | pICE_Low_copy_bb | CATTACTC |
| BB_components | pL0R-lacZ-4_bb | cattactgc |
| BB_components | pSa | GATCCCC |
| Selectable Markers | SpecR | atgggggaa |
| Selectable Markers | KanR | Ctaaaacaa |
| Selectable Markers | nptI | TTAGAAA |

[Supplementary Table 4] A complete list of sequences used.
